# Diversity and abundance of archaeal amoA genes in the permanent and temporary oxygen minimum zones of Indian Ocean

**DOI:** 10.1101/2022.04.27.489739

**Authors:** Prasannakumar Chinnamani, Anandjothi Elamaran

## Abstract

Oxygen minimum zones are results of oxygen consumption exceeding the oxygen availability in stratified water columns of the marine environment. We compared the ammonia monooxygenase subunit A (amoA) gene abundance and the diversity of ammonia-oxidising archaea (AOA) in the Arabian Sea (AS) with those of the Bay of Bengal (BoB). Three primer pairs targeting amoA genes of water column A (WCA), water column B (WCB) and total AOA (amoAt) captured different densities of gene copy numbers in both marginal seas. Water column A (WCA) ecotypes were more abundant in the AS than in the BoB. Core-OMZ depths of the BoB contained 10 times lower amoA copy numbers than those of the AS. Along with sampling depth, concentration of ammonia shapes the WCA/WCB ecotypes in AS/BoB. Among the total AOA populations, WCB ecotypes were more abundant. The amoA gene sequences were either of Nitrosopumilales or Ca. Nitrosotaleales members and belonged to NP-γ, NP-δ, NS-β, NS-γ and NS-ε sub-clades. Pairwise distance and nucleotide diversity index analysis reveals that BoB nurtures two times more diverse amoA sequences than the AS. The core OMZ region of the BoB contains a two-fold higher diversity of amoA gene sequences compared to the AS, whereas the AS contains 13 times more abundant amoA copies than the BoB.

## INTRODUCTION

In stratified water columns of marine environments, when the demand for oxygen exceeds oxygen availability during organic matter decomposition, oxygen minimum zones (OMZs) are formed, in particular, when the oxygen concentration drops below 20 µM (Paulmier and Ruiz-Pino, 2009). Such zones are currently expanding due to global warming (Whitney et al., 2007; Falkowski et al., 2011), and active climate gasses such as carbon dioxide (CO_2_), nitrous oxide (N_2_O) and methane (CH_4_) are produced during microbial organic matter degradation in OMZs (Paulmier and Ruiz-Pino, 2009). Currently, OMZs occupy about 7% of the ocean volume (Paulmier and Ruiz-Pino, 2009; Wright et al., 2012). In OMZs, ammonia-oxidizing archaea (AOA) along with *Nitrospina, Planktomycetes* and SUP05 have been recognised as key players of the carbon, nitrogen and sulphur cycles (Hawley et al., 2014; Hallam et al., 2006; Walsh et al., 2009; Walker et al., 2010). Being ubiquitous and abundant in the marine environment, AOA are major players of nitrification (conversion of ammonia to nitrate) (Stahl and Torre, 2012) and significant contributors to carbon fixation processes (Hansman et al., 2009). They also play a role in methane production (Metcalf et al., 2012) and cobalamin synthesis (Heal et al., 2016). The amoA genes that encode for ammonia monooxygenase subunit A have been sequenced for studying the diversity and ecology of AOA (Zheng et al., 2017). After the 16S ribosomal RNA (rRNA) gene, amoA is the second-largest sequenced gene in microbial ecology, comprising 56% of the sequences available in GenBank (∼68,000 archaeal sequences as of November 2019, excluding short fragments from high-throughput sequencing).

The abundance of archaeal amoA genes is one to three orders higher than that of the bacterial amoA gene in several oceans (Wuchter et al., 2006a; Mincer et al., 2007; Agogue et al., 2008; Beman et al., 2008). The abundances of amoA and 16S rRNA genes of Thaumarchaeota in the Arabian Sea (AS) reveal that the majority of Thaumarchaeota were AOA (Pitcher et al., 2011), despite the fact that amoA genes per cell could vary based on environmental parameters (Wuchter et al., 2006; Agogue et al., 2008). Also, AOA amoA or 16S rRNA gene copies and rates of ammonia oxidation are positively correlated, revealing their dominant role in ammonia oxidation in the coastal eastern Pacific (Santoro et al., 2010), the Gulf of California (Beman et al., 2008) and the North Sea (Wuchter et al., 2006a). In addition, similar patterns of abundances did not correlate with oxidation rates in OMZs of AS (Newell et al., 2011), the Central California Current (Santoro et al., 2010) and the South Pacific Ocean (Lam et al., 2009).

Interestingly, the obligate aerobic AoA amoA gene copy numbers are abundant in OMZs, but it is unclear which metabolic function supports their survival and growth in OMZs of AS. This may be due to the micro-molar concentration of oxygen or the ammonium input from the mesopelagic waters below the OMZs. Despite their close geographical locations, AS and BoB are highly different. The BoB is less productive than the AS, as the flow of nutrient-rich water into subsurface and oxycline layers of the AS makes them highly productive, while this process is unlikely in the BoB (Kumar et al., 2002). The OMZ of the Arabian Sea is strong, whereas that of the Bay of Bengal is relatively weak (Ittekkot et al., 1991; Rao et al., 1994; Naqvi et al., 1994); the condition in the BoB is possibly due to the robust stratification and weak upwelling (MsCreary et al., 2013). While nitrate is intensely being removed in the nearby OMZ of the AS, such volumes of loss do not occur in the OMZ of the BoB, which could be attributed to the differences in the rates of decomposition of exported organic matter throughout the water column (Azhar et al., 2017). Another study confirmed that the abundant availability of nitrate and the highly variable oxygen concentrations inhibit nitrate loss in the BoB (Johnson et al., 2019).

Numerous amoA-based studies proved that AOA diversity and abundance depend on multiple factors and are strongly partitioned by ecosystem (Francis et al., 2005; Biller et al., 2012; Cao et al., 2013; Yao et al., 2013; Sintes et al., 2013; Restrepo-Ortiz et al., 2014). Although differences in abundance and diversity of AOA were clearly observed among geographically distinct seas, geochemically similar seas may also house distinct AOA populations (Techtman et al., 2017). Thus, biogeography only in part contributes to determining the diversity and distribution of AOA. Thus, true drivers of AOA diversity and distribution are still unknown, especially in the global OMZs. Peng *et al*. (2013) conducted a comparison between AS (surface and anoxic layers) and the South Pacific Ocean and showed that the AOA community composition was determined by the topography and communication of AOA and anammox bacteria in their respective seas. We therefore hypothesise that the differences in mineralisation depth (Azhar et al., 2017) and variable oxygen concentrations in the BoB and the AS (Johnson et al., 2019) could have shaped unique OMZ features, which in turn could be shaping specific amoA phylotypes in the respective seas.

## METHODS

### Sample collection and physicochemical parameters

Six sampling sites were chosen in the Arabian Sea and the Bay of Bengal, and water samples were collected during the post-monsoon season of November 2011 and January 2012 on FORV Sagar Sampada. The latitude and longitude positions of the sampling sites are provided in Table 1. We selected three sampling stations in the Arabian Sea (1, 2 and 3) and three in the Bay of Bengal (5, 6 and 7). Maximum water depth ranged between 1,700 and 4,400 (m) for all stations (Fig. 1). All three stations in the Arabian Sea are within a well-documented OMZ with abundant AOA (Newell et al., 2011; Pitcher et al., 2011). In contrast, stations 5 and 6 of the BoB are within a well-documented OMZ with AOA (Bristow et al., 2017). The oxygen concentration was < 10 µM in the core the OMZ of both seas (Fig. 2). Samples were collected in 12-L Niskin bottles (12) mounted on a conductivity-temperature-depth (CTD) rosette system (SeaBird Electronics). Physico-chemical parameters of seawater (conductivity, density, depth, dissolved oxygen (DO), pH, turbidity, salinity and temperature) were recorded with appropriate sensors attached to the CTD system.

**Fig. 1:**
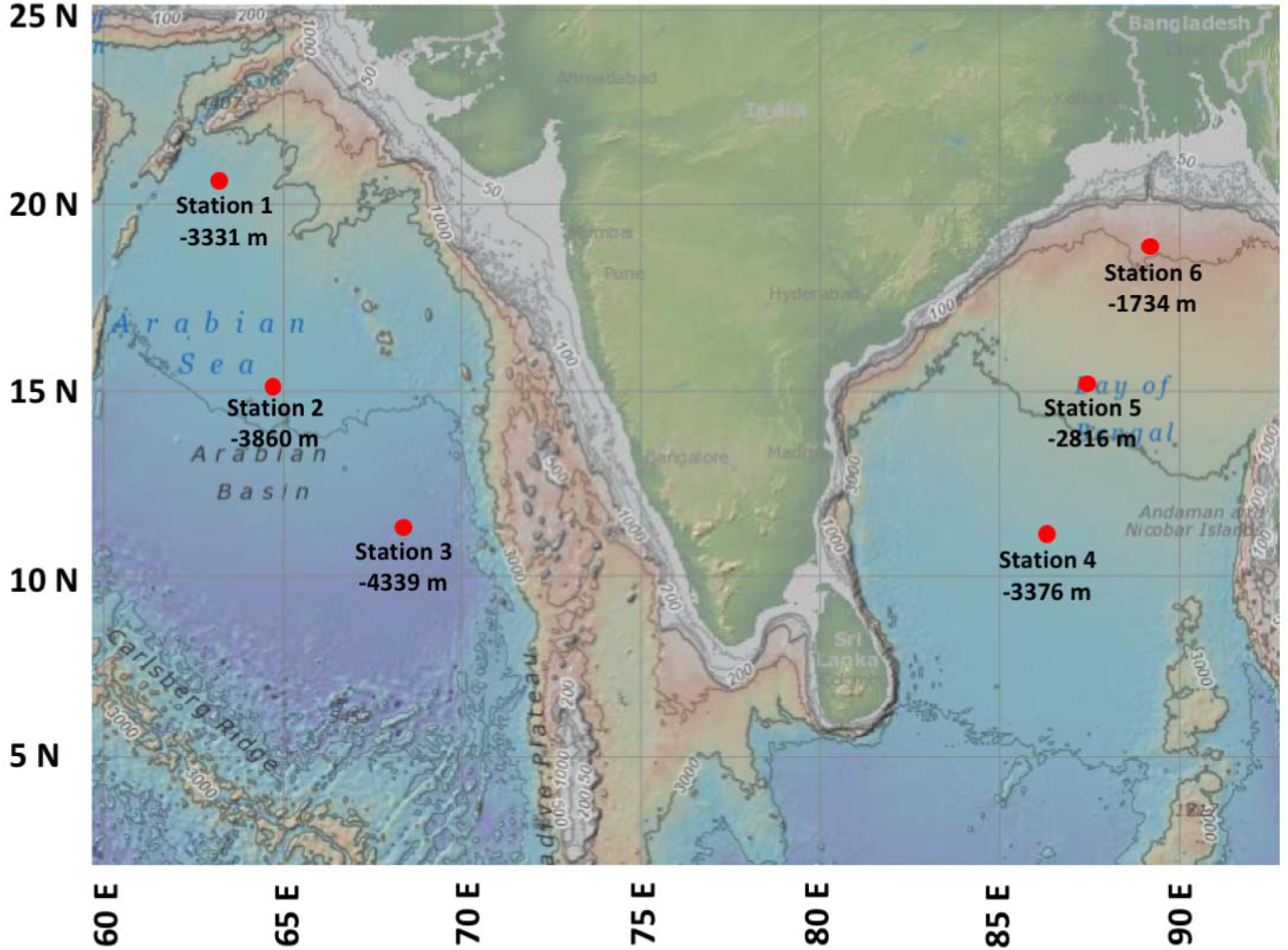
Bathymetry map of sampling stations with maximum depths (in meters) in the Arabian Sea and the Bay of Bengal

**Fig. 2:**
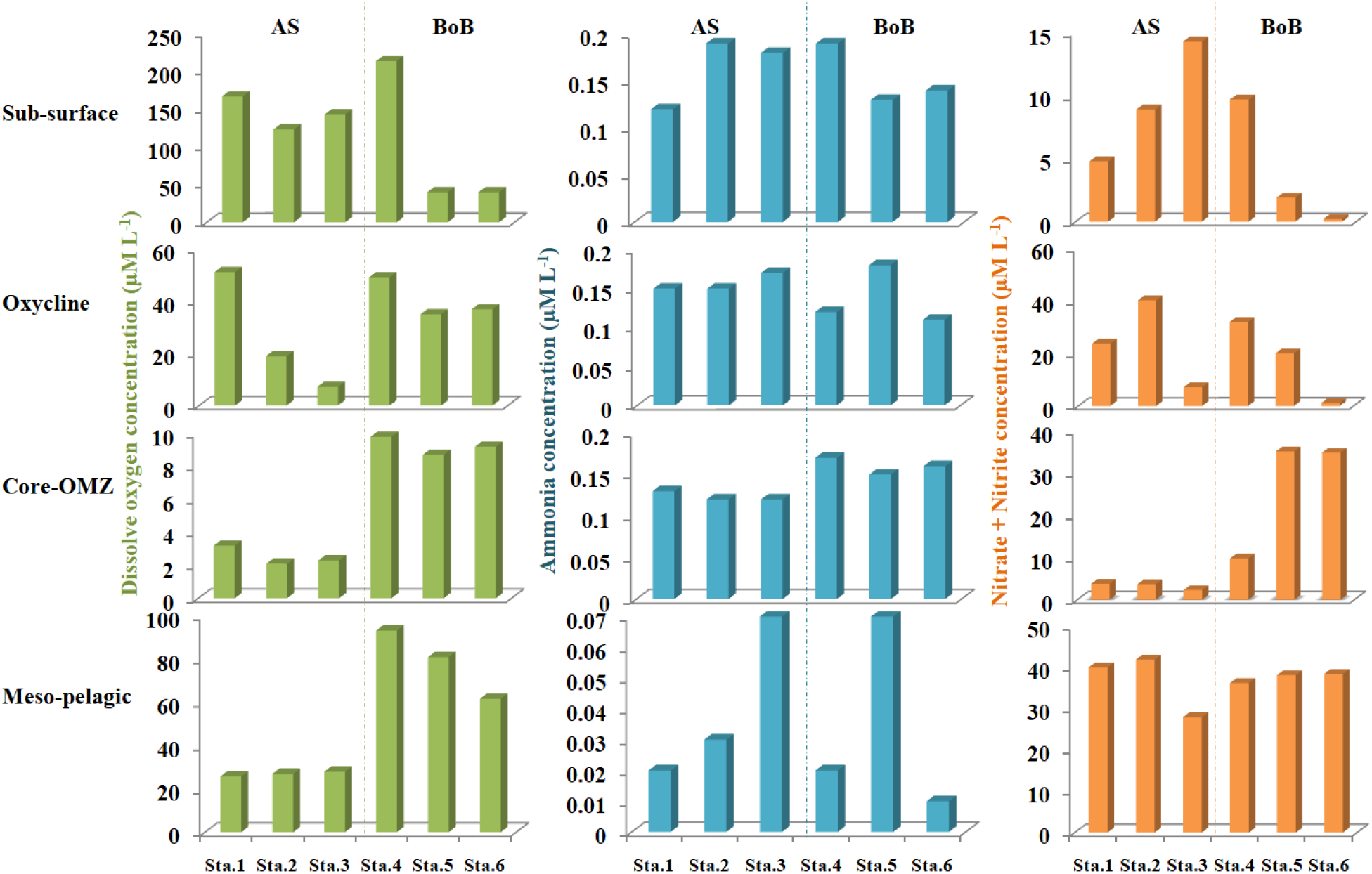
Depth profiles of dissolved oxygen, nitrite + nitrate and ammonia concentrations at the six sampled stations. Station numbers are indicated by numbers 1-6, and sampled depths are indicated by ‘a to d’.

Sampling depths were fixed based on dissolved oxygen (DO) data obtained during CTD (Table 1) deployment. Water samples for gene abundance were collected from four different depths at each station; a) subsurface (∼20 m in all stations), b) oxycline waters (110–140 m in AS; 140–210 m in BoB), c) the core of the OMZ (350-390 m in AS; 600-710 m in BoB) and d) mesopelagic waters (O_2_ conc. > 10 uM) (> 1,500 m in AS; > 900 m in BoB). The station names were indicated by station number (1 to 6), followed by sampled depths (a to d), i.e. ‘4.d’ indicates mesopelagic water of station 4. The DNA samples (triplicates) for diversity assessment were collected from the core OMZ of the stations 1 and 6. A nutrient AutoAnalyser (*QUAATRO;* Bran+Luebbe) was used to measure dissolved concentrations of nitrate (NO_3_-N), nitrite (NO_2_-N) and ammonium (NH_4_-N).

### DNA extraction, qPCR, PCR, cloning and sequencing

Approximately 20 L of seawater were filtered through 0.22-μm polycarbonate membrane filter paper (47 mm diameter; Millipore), and the polycarbonate membranes were flash-frozen in liquid nitrogen and stored at -80°C until further analysis. The DNA was extracted using the DNeasy kit (Bioserve, India) following the manufacturer’s protocols. Concentration and purity of the genomic DNA were checked with a NanoDrop spectrophotometer (Thermo Scientific 2000/2000c) (Johnson, 1994). The DNA yields ranged from 0.08 to 3.15 µgL^-1^ of seawater across all samples.

Abundance of the archaeal amoA genes was determined using three sets of primers, targeting 1. total archaeal amoA (amoAt); Arch-amoAF/Arch-amoAR (Francis et al., 2005), 2. water column A (WCA) ecotypes; Arch-amoAFA & Arch-amoAR (Mosier and Francis 2011) and 3. water column B (WCB) ecotypes; Arch-amoAFB& Arch-amoAR (Mosier and Francis 2011). The QPCR quantification was carried out as follows: initial denaturation at 95°C for 10 min, followed by 40 cycles of 95°C for 15s and primer annealing at 56°C for 1 min, followed by detection. All reactions were carried out in triplicate. In this study, for the water column C, the soil assemblage primer set was not used because in marine systems, amoA gene (WCC) abundance is low (Francis et al., 2005). For cloning and sequencing, linearized plasmid vectors were used (TOPO vector, Invitrogen); the plasmids were linearized by the *NotI* restriction enzyme (DNeasy clean kit, Bioserve, India) and stored at -80°C until future use. The vectors were freshly diluted prior to the experiments. The WCA and WCB ecotypes were quantified with identical reaction chemistries as follows: 12.5 µL Taqman Master Mix 2.0 (Bioserve, India), 200 nM of each primer, 300 nM of each probe and 1 µL DNA template per reaction, with a final volume of 25 µL. Efficiencies for all qPCR assays ranged from 92 to 99% across all samples.

For DNA sequencing, replicate DNA extractions (from stations 1 and 6) from three sub-samples were pooled from each sampling and extracted using the Power Water DNA isolation kit (MOBio, USA). Archaeal amoA fragments (∼635 bp) were amplified using the primers Arch-amoAF and Arch-amoAR under standard PCR conditions (Beman& Francis, 2006). Each sample was amplified thrice and pooled to minimise PCR bias; the products were gel-purified and ligated into pMD19 vectors (Bioserve Biotechnologies, pvt.ltd., India) according to the manufacturer’s instructions. *Escherichia coli* TOP10-competent cells were used for hybrid vector transformation (Sambrook& Russell, 2001), and recombinants were selected by using X-Gal-IPTG in Luria–Bertani (LB) indicator plates supplemented with 100 mg ampicillin ml^-1^. Approximately 390 white colonies were randomly selected from each clone library, and cloned amoA fragments were re-amplified using the primer pairs M13-D (5’-AGGGTTTTCCCAGTCACGACG-3’) and RV-M (5’-GAGCGGATAACAATTTCACACAGG-3’). A later primer pair (RV-M) was used for sequencing with an ABI 3770 automatic sequencer (Applied BioSystems) at Bioserve Biotechnologies, Pvt. Ltd. (India).

### DNA sequence analysis: CHIMERA check, phylogeny

The DNA sequences were extracted using ChromasPro (ver. 1.9.9), and the total sequences were subjected to the *chimera* filter using UCHIME (Edgar et al., 2011). Resultant non-chimeric sequences were grouped into operational taxonomic units (OTUs) based on a cut-off value of 95% (Christman et al., 2011). The DNA sequence similarities between two sampling regions were calculated with Libcompare of the DOTUR program (Schloss & Handelsman, 2005) and aligned using the CLUSTAL_X program (Thompson et al., 1994). We used total amoA sequences representing global amoA phylogeny, classified and annotated by Alves et al. (2018), as a reference file, as it represents a complete picture of amoA phylogeny to date. The phylogenetic tree was constructed using MEGA X (Kumar et al., 2018) and refined usingiTOL (Letunic and Bork, 2019).

### Statistical analysis

Pairwise distances and nucleotide diversity were calculated within marginal sea sequences using Kimura-2 parameters in MEGA X (Kumar et al., 2018). The sequence coverage of the archaeal amoA gene was estimated using a rarefaction curve plotted in ‘Fungene’, a functional gene pipeline and repository in the Ribosomal Database Project (RDP) (Cole et al., 2013). The statistical inference helps to compare the complexities between two or more communities and the sampling efficiency. Correlations between AOA abundances and environmental variable in all sampled stations were analysed by canonical correspondence analysis (CCA) using PAST (Hammer et al., 2001). Shannon diversity index, evenness and sampling station similarities were plotted in PAST.

## RESULTS

### Physiochemical properties

The bathymetry of sampling stations in AS and BoB is provided in Figure 1. Overall, the average surface temperature was 25°C in both the marginal seas, whereas the oxycline region showed a different temperature; i.e. the average temperature of the BoB (25°C) was slightly higher than that of the AS (19.8°C) (Table S1). Overall, the temperature gradient between oxycline and the core of the OMZ was relatively higher in the BoB (∼25 and 13.5°C) than in the AS (∼19.8 and 11.5°C) (Table S1). Among the sampling stations, BoB was ∼5, 2 and 5°C warmer than the AS in oxycline, core OMZ and mesopelagic waters, respectively. Sampling depth ranged from subsurface (∼20 m; in all stations) to a maximum of 1,620 m (at mesopelagic depths of station 3 in AS).

The core OMZ of the AS was three times stronger (with ∼3 µM DL^-1^) than that of the BoB (∼9 µM DL^-1^), while nitrite accumulation was 9-fold higher in the BoB (up to 35 µM) than in the AS (up to 3.2 µM) (Fig. 2). The core OMZ of the BoB had a higher ammonia concentration (0.16 µm N L^-1^) than that of the AS (0.12 µm N L^-1^) Overall, the subsurface stations of the AS were better oxygenated than that of the BoB, whereas mesopelagic water of the BoB was better oxygenated than that of the AS. Regarding ammonia concentrations, a maximum of 0.2 µm N L^-1^ was recorded in subsurface depths of station 2 (AS) and station 4 (BoB). All mesopelagic depths in both marginal seas had minimum ammonia concentrations, with the lowest being observed for station 6 (0.01 µm N L^-1^) in the BoB. (Table S1).

### Variations in AoA amoA gene abundances in AS and BoB

The WCA and WCB ecotypes were detected in all stations and were more abundant in the AS (in the order of 10^6^ and 10^7^ gene copies L^-1^, respectively) than in the BoB (in the order of 10^4^ and 10^6^ gene copies L^-1^, respectively) (Fig. 3, Table S2). In AS stations, a high abundance of WCA ecotypes (1.9 × 10^7^ copies) was observed in the core OMZ (station 1), whereas a minimum of 22 copies were recorded in mesopelagic depths of the same station. In the case of BoB stations, a maximum of 2.7 × 10^5^ gene copies of WCA was recorded in subsurface waters of station 6 and a minimum of near-detection limit (7 copies L^-1^) was recorded in mesopelagic waters of station 5. However, in both marginal seas, amoAt copy numbers were in the order of 10^7^ copies L^-1^, indicating that the two marginal seas contain similar densities of AOA but of a different species composition, as the WCA/WCB primer used picked up different densities of amoA copy numbers at different depths among the stations (Fig. 3, Table S2). Among all stations, the abundance trend of amoAt copy numbers followed the order subsurface < oxycline > core OMZ < meso-pelagic waters. In all subsurface stations of the AS, WCA ecotype amoA copy numbers were high (∼10^5^), and the numbers decreased in the oxycline layers, followed by a sharp increase to reach the maximum in the core OMZ (∼10^6^) and a sharp decline in the mesopelagic depths (10^2^). Generally, the WCB ecotypes were lowest in subsurface waters and highest in the core OMZ of the AS (10^7^ copies L^-1^), followed by a decline (10-fold) in mesopelagic depths. Such observational trends were common globally, but such trends were steeper in BoB stations where the decline is 100 fold lesser in mesopelagic depths.

**Fig. 3.**
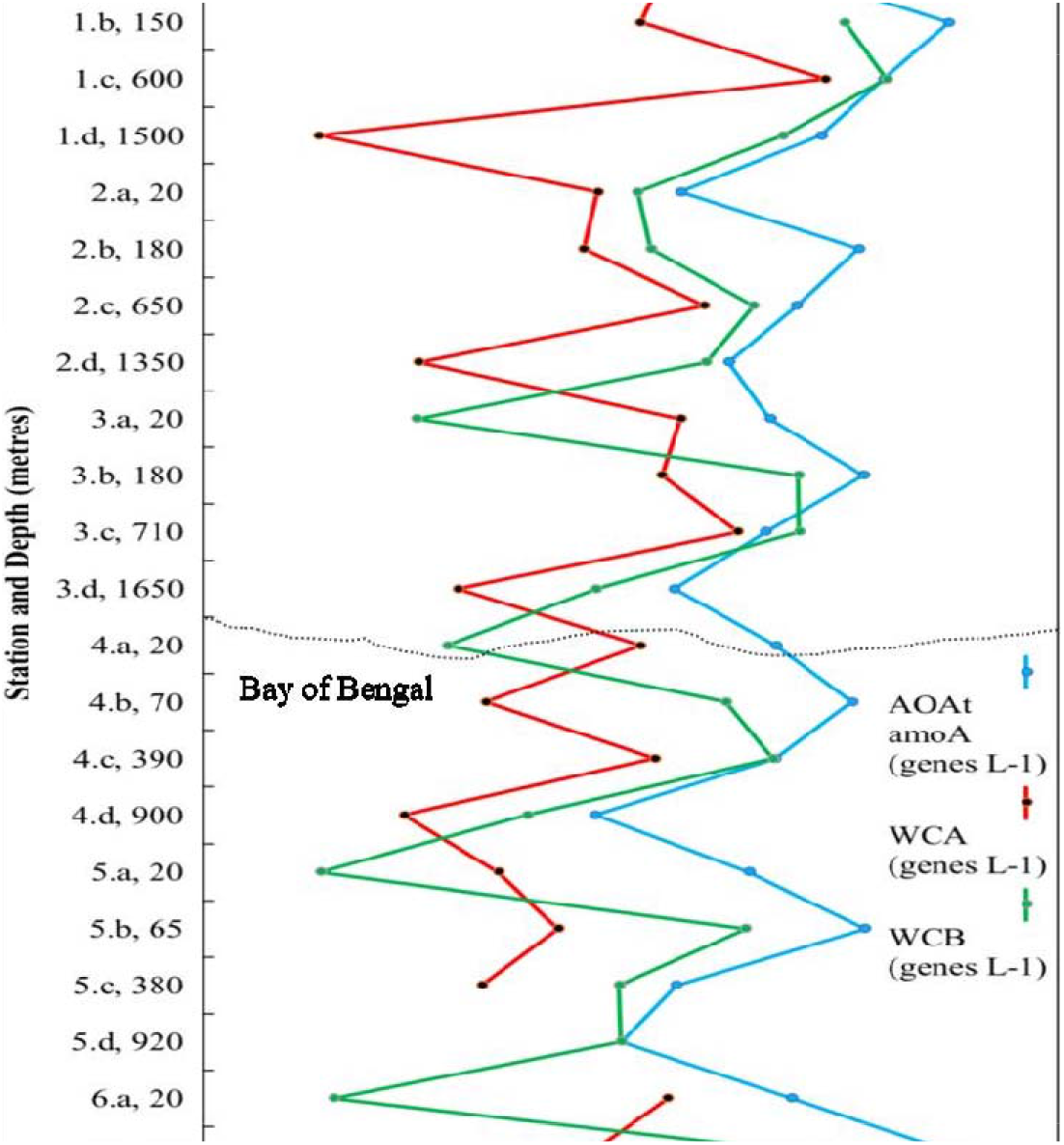
Abundance of total archaeal amoA (amoAt), water column A (WCA) and water column B (WCB) ecotypes. Sampled stations (1, 2, 3, 4, 5, 6), depth profile (a = surface waters; b = oxycline waters; c = core oxygen minimum waters and d = mesopelagic waters) along with depth (in meters) are provided on the x axis and amoA gene copy numbers on the y axis.

### AOA amoA abundance in core OMZs (AS vs. BoB)

The AoA amoAt and WCB-amoA gene copies clearly dominated throughout the core OMZ (Fig. 4). The WCA-amoA, WCB-amoA and amoAt gene copies were 90, 23 and 7.5 times higher in the AS than in the BoB (Fig. S3, Table S2).

**Fig. 4:**
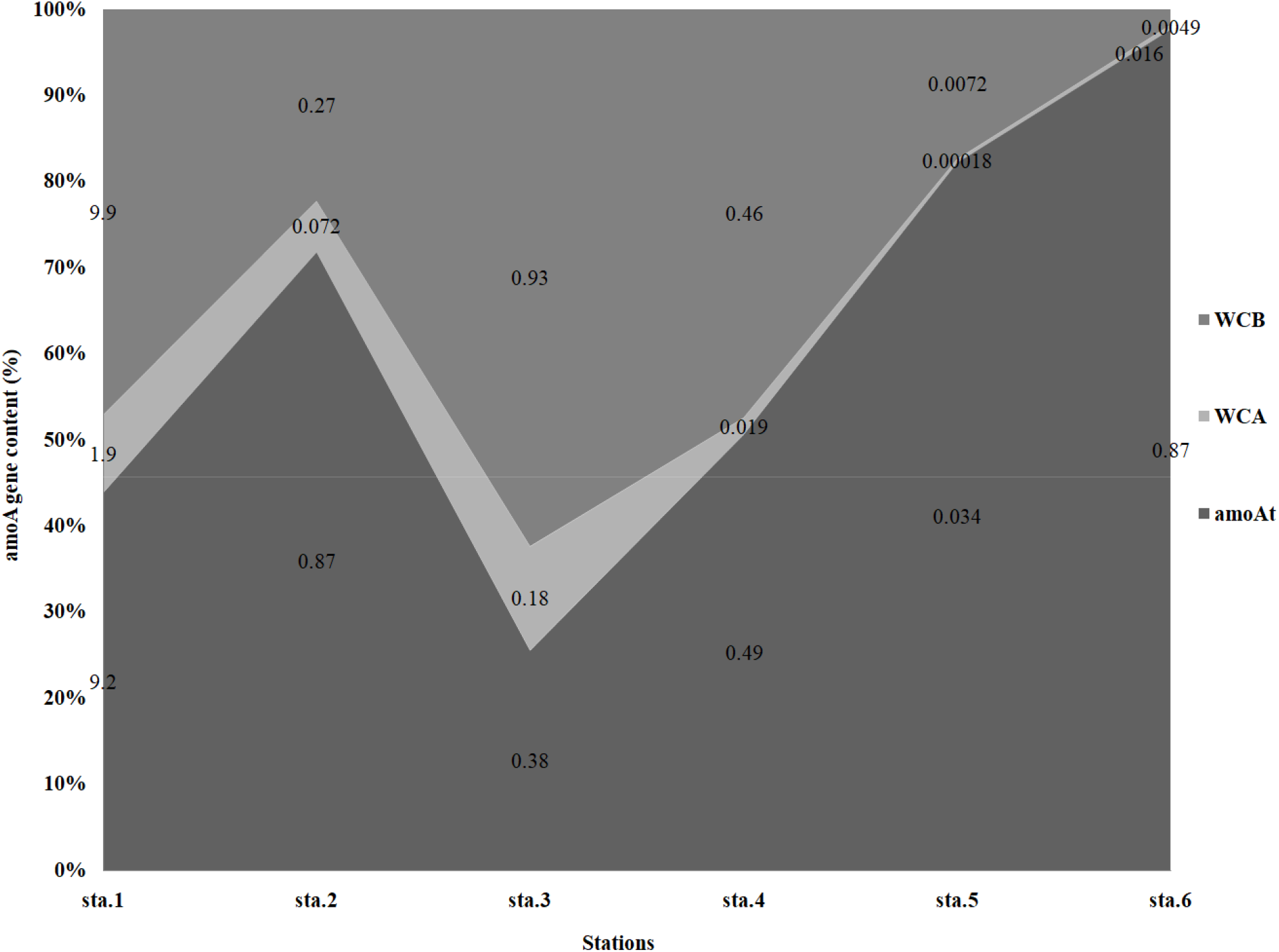
Percentage compositions and abundances of amoAt, WCA and WCB amoA gene copies in the core OMZ of the AS (sta. 1, 2 & 3) and the BoB (sta. 4, 5 & 6). Numbers inside the plot represent gene copy numbers (x 107) per litre of the samples of the corresponding stations.

Among the stations, WCA-amoA, WCB-amoA and amoAt were more abundant in the AS station 1(9.2, 1.9 and 9.9 (× 10^7^ copies L^-1^), respectively) than in the BoB station 6 (8.7 × 10^6^ copies L^-1^), justifying the selection of these stations (sta. 1 & sta. 6) for sequencing studies. Surprisingly, WCB amoA copies outnumbered amoAt copies in station 1 (9.9 × 10^7^ copies L^-1^) (Fig. S3), leading us to infer question whether the actual value represents a true phenomenon in a sampled station, qPCR contamination during experimentation or the inability of the used amoAt primer to capture entire AOA amoA gene copies. Also, in station 1, we recorded a higher abundance of WCA-amoA gene copies (1.9 × 10^7^ copies L^-1^) when compared to the other stations. Regarding the WCA, WCB and amoAt copy numbers, the AS contained 13 times more amoA copies than the BoB

### Influences of environmental parameters on WCA, WCB and amoAt gene copies

The WCA amoA abundance and ammonia concentration were positively corelated, while WCA was partially negatively correlated with sampling depth (Fig. S2), We also found a partial negative correlation between abundance of WCB ecotypes and dissolved oxygen concentration, explaining its relatively lower abundances in mesopelagic waters below the OMZs of both marginal Seas (Fig. 5, S2); amoAt abundance was positively correlated with WCA ecotype.

**Fig. 5:**
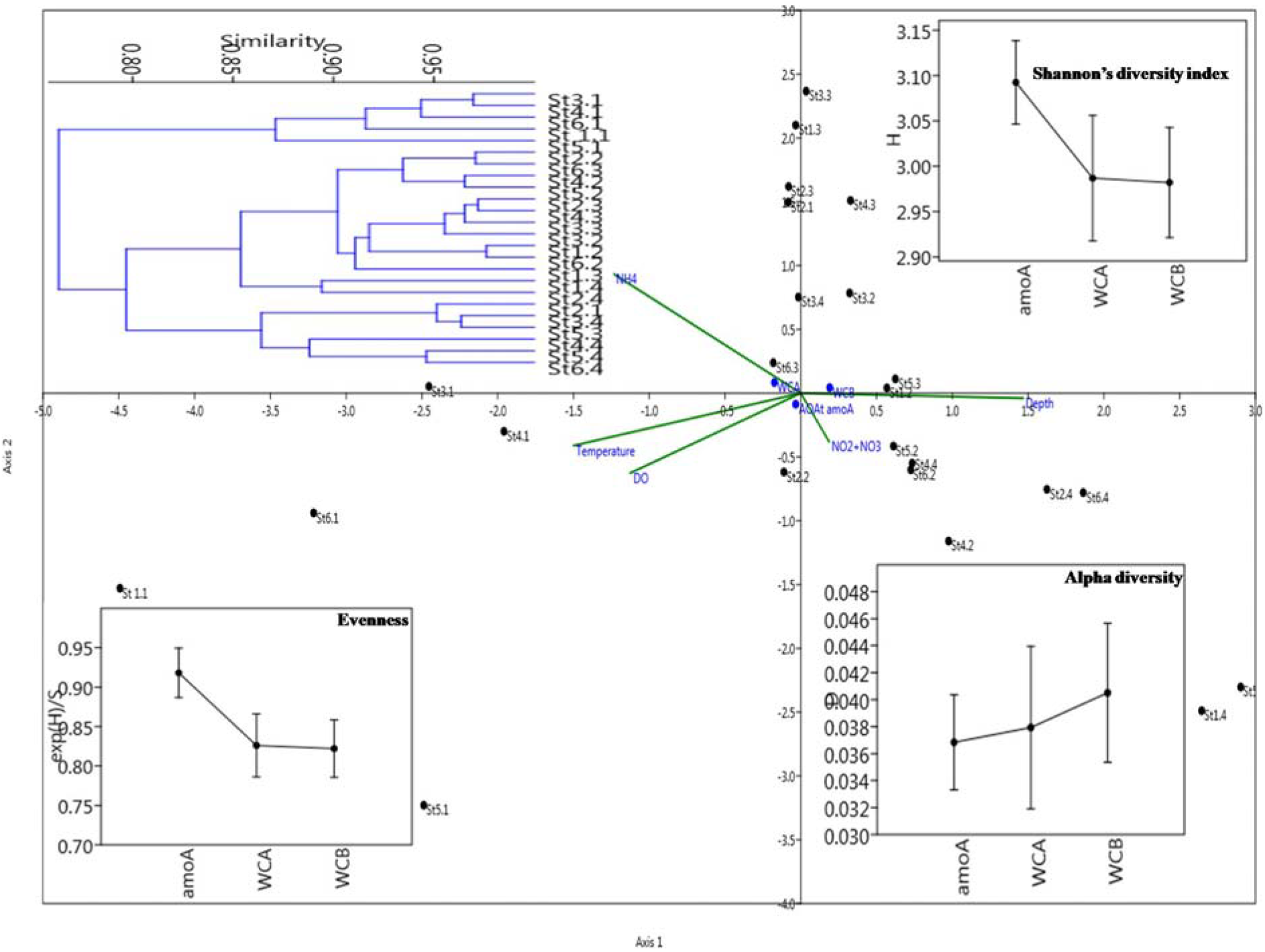
CCA plot describing relationships between physicochemical and biological parameters within sampling stations, along with diversity indices.

Not more than 40% variations were explained by combining both axes in the CCA plot, indicating that factors responsible for shaping the community variation were not included in the present study (Fig. 5). Among the environmental and biological parameters, subsurface samples, station 3 (AS) and station 4 (BoB) were highly (> 95%) similar (Fig. 5). Among oxycline samples, station 1 (AS) and 6 (BoB) and stations 4 and 5 (BoB) were highly similar. Regarding the OMZ layers, station 2 (AS) and 4 (BoB) were relatively more similar to each other compared with station 3 (AS) (Fig. 5). Overall, amoAt showed higher Shannon diversity and evenness values in the stations, and alpha diversity followed the order WCB > WCA > amot (Fig. 5).

### AOA phylogenetic diversity

We sequenced 390 amoA genes from the AS and the BoB, of which 22 sequences were chimeric (5.6% of total sequences); 190 amoA (AS) and 178 (BoB) sequences were included in further analyses. Clustering analysis grouped the sequences into 45 OTUs, of which 19 were from the AS and 26 from the BoB. The phylogram constructed using the global reference file of Alves et al. (2018) and the representative OUT sequences from the present study revealed that all OTUs belong to two main classes, viz; Nitrosopumilales (NP) and Ca. Nitrosotaleales (NT) (Fig. 6). Members of NP were segregated into alpha (NP-α), gamma (NP-γ), delta (NP-δ) and epsilon (NP-ε) clades, whereas the members of NS were segregated into alpha (NS-α), beta (NS-β), gamma (NS-γ) and epsilon (NS-ε) clades. Sequences from the BoB had representatives in all clades, whereas sequences of the AS were represented only in NP-α, NP-γ and NP-δ clades.

**Fig. 6:**
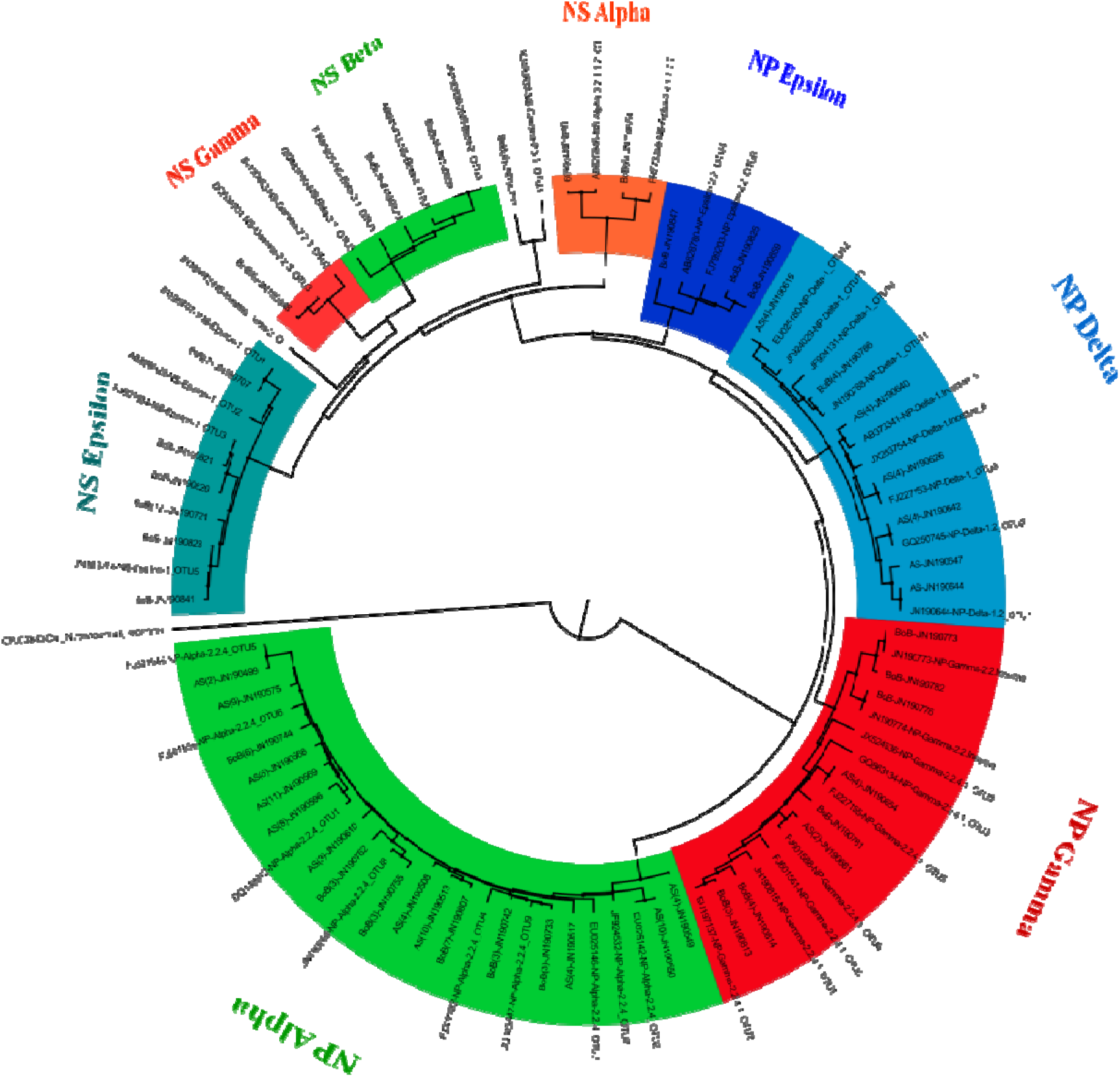
AoA amoA phylogeny retrieved from the core OMZ of the Arabian Sea and the the Bay of Bengal.

Even with limited representation in NP clades, amoA sequences of AS constituted 65 and 85% of total OTUs in NP-α and NP-δ clades, respectively. Even from the global reference file of Alves et al. (2018), one representative OUT in the NS-γ clade could not be grouped (Fig. 6). We found that Alves et al. (2018) have identified seven of the sequences (published in GenBank) from the present study to define the global phylogeny, consisting of clades NP-γ-2.2_Incertae, NP-γ-2.2.4.1_OTU1, NP-δ-1_OTU11, NP-δ-1.2_OTU1, NS-β-2_OTU3, NS-γ-2.2.3_OTU2 and NS-ε-1_OTU1 (Fig. 6). The respective GenBank accession numbers of those sequences were JN190774, JN190815, JN190788, JN190644, JN190707, JN190852 and JN190828, respectively. Among the unique representatives of global phylogeny identified in the present study, amoA sequences of the AS were represented only in the NP-δ-1.2_OTU1 clade, whereas the remaining ones (six) were occupied by BoB amoA sequences.

The overall pair-wise nucleotide distance data revealed that amoA sequences of the BoB (0.28) were 2.3 times more abundant than those of the AS (0.12), which is surprising given the overall dominance of AS in amoA gene copies. The nucleotide diversity index (π) of the BoB (0.23) was two times higher than that of the AS (0.11). Overall, the core OMZ region of the BoB contained twofold diverse amoA gene copies compared to that of the AS, posing the question whether a weaker OMZ promotes a higher AOA species diversity (based on the BoB scenario) or whether reduced oxygen levels reduce the Thaumarcheota population? (based on the AS scenario). A rarefaction curve as generated to analyse the depth of the sampling site, indicating cut-off at 95% (Fig. S1). The BLAST analysis revealed that all sequences were matched with unculturable amoA sequences produced elsewhere from diverse environments, including marine and coastal/wetlands (Table S4). The sequences generated from the present study were added to GenBank under the accession numbers JN190498-JN190865.

## DISCUSSION

Sampling two marine systems consecutively in the same season for comparison of biological data was a difficult task and required considerable planning. The *amoA* gene has been extensively used to estimate the abundance and diversity of AOA and has provided evidence for their high phylogenetic diversity in natural environments (Prosser and Nicol, 2012). The present study documented amoA abundances and diversity in AS and BoB in the same cruise track. Among the stations in the AS, amoA abundances were one-to two-fold higher compared to the values found in a previous study in nearby sampling sites (Newell et al., 2011) and in the Atlantic Ocean (Wuchter et al., 2006).

The amoA copies of WCA, WCB and amoAt were detected in all depths and stations, similar to observations made in North Eastern Pacific waters (Smith et al., 2015). However, unlike in North Pacific waters, where amoA abundances ranged between 10^3^ and 10^6^ copies per litre, AS/BoB contained highly variable abundances among the sampling stations (amoAt copies between 10^1^ and 10^8^ copies per litre), which may be due to the geographically larger sampling area and the multiple depth profiles (four depths) per station. In this study, thehe patterns of high abundance of amoAt copies, concentrated from oxycline to core OMZ depths and decreasing in mesopelagic waters, resembles with the patterns of North Pacific waters (Smith et al., 2015).

Also, the patterns of WCA, being one order less abundant in both marginal seas, and of WCB, being highly variable along the depths, were similar to North Pacific waters (Smith et al., 2015). Our data support the previous hypothesis that WCA can adhere to relatively higher concentrations of ammonia than WCB. This is also supported by a positive correlation between WCA and ammonia concentration. Apart from depth, ammonia concentration appears to shape the WCA/WCB ecotypes in AS/BoB, as the variable concentrations of ammonia drive AOA diversity (Sintes et al., 2013). Also, the positive correlation observed between WCB and amoAt copy numbers in both marginal seas implies that WCB could be a dominant community among the total AOA populations (especially in station 1 of the AS) (Hansman et al., 2009).

The 16S rRNA and *amoA* gene of AOA were interchangeably used in phylogenies, as both the topologies of the trees were similar with higher taxonomic resolution (Nicol et al., 2008). To assess environmental patterns and global AOA distribution, Alves et al. (2018) used 33,378 amoA sequences of AOA and found that the global phylogenetic distribution of AOA contains four major clades, viz., 1. Ca. Nitrosocaldales (NC), Nitrososphaerales (NS), Ca. Nitrosotaleales (NT), Nitrosopumilales (NP) and a minor clade named NT/NP-Incertaesedis (NT/NPIS). These clades, grouped in the level of classes, were further subdivided into sub-clades as alpha, beta, gamma, etc., and the sub-clades were furthermore divided based on a number of representative OTUs in the group (1.1, 1.2, etc.). All amoA sequences from both marginal seas represented one group in NP (77.8%) and NS (22.2%). Members of NP were segregated into alpha (NP-α), gamma (NP-γ), delta (NP-δ) and epsilon (NP-ε) clades, whereas the members of NS were segregated into alpha (NS-α), beta (NS-β), gamma (NS-γ) and epsilon (NS-ε) clades.

Spurious AOA from lineage NS have also been found in marine sediments (Alves et al., 2018); in the present study, one representative OTU in NS-γ clade did not group together. Globally, 84% of NS-γ are most specifically associated with soils-sediments (Alves et al., 2018). Hence, regarding the origin of representative OTUs in the present study, it is unclear whether they represent thriving organisms or dormant/dead cells transported from the land.

Although NP-α and NP-ε OTUs were dominant among the NP clade, globally, only 45% of lineage NP were actually detected in unambiguously marine environments (Alves et al., 2018). Overall, the majority of NS and NP members represented from the AS and the BoB was globally recognised to occur in marine systems, among which the AOA of the BoB were more diverse than those of the AS.

Using the sequences described in Alves et al., (2018), we found that seven OTUs were represented in global phylogeny, where the AOA of the BoB contributed six different OTUs. These seven OTUs were clade representatives, viz.; NP-γ-2.2_Incertae, NP-γ-2.2.4.1_OTU1, NP-δ-1_OTU11, NP-δ-1.2_OTU1, NS-β-2_OTU3, NS-γ-2.2.3_OTU2 and NS-ε-1_OTU1BoB were unique OTUs, which has been recovered by stringent chimeric screening and multiple re-drawn phylograms (Alves et al., 2018).

Another interesting phenomenon is the absence of AS amoA sequences in the NS clade. Further studies should therefore investigate the drivers of amoA diversity in both marginal seas, as they seem to incubate different amoA populations. Even though the factors driving these OMZ-specific amoA communities were unclear in the present study, when comparing the AS with the BoB, containing nano-molar concentrations of O_2_ in the OMZ, which inhibit stable accumulation of nitrite (Bristow et al., 2016), differences in adiabatic factors (Prasanna Kumat et al., 2002) cannot be ruled out.

Peng et al. (2013) conclude that the AOA community in OMZs of the AS is significantly different from that of the Eastern Tropical South Pacific, and such differences were due to geographic factors rather than environmental parameters. Although our results do not agree with previous findings, as AOA amoA populations in geographically closer OMZs are genetically distinct, differences in the depths of the OMZs in both marginal seas could be a potential factor. Also, the subsurface temperature in the BoB was higher than that in the AS, and according to previous findings, a 1.5-2°C higher temperature in the BoB compared to the AS leads to strong stratified surface layer in the BoB, limiting the mixing of nutrients from below the surface layer (Prasanna Kumar et al., 2002). However, in changing OMZ boundaries of global ocean (Stramma et al., 2011), the threshold of resistance of the BoB is unknown.

## Conclusions

Although Sangar’s sequencing has a high power in amplicon length recovery, thereby revealing more information per DNA sequence, the quantity of data provided by Illumina sequencing is unmatchable. As unique clades were discovered by Sangar’s sequencing in the present study, further studies with next-gen sequencing platforms are needed to investigate the differences between the AS and the BoB in terms of the AOA population. The AOA of the BoB contained twofold diverse amoA gene copies compared to the AS, posing the questions whether a weaker OMZ promotes a richer AOA diversity or whether lower oxygen levels reduce the Thaumarcheota population.

## Acknowledgement

Authors thank the Department of Biotechnology, Ministry of Science and Technology, Government of India for the financial assistance provided through INSPIRE fellowship (IF10431) and China Postdoctoral Science Foundation, China, for financial assistance provided through National Postdoctoral Fellowship (Ref.: 0050-K83008).

## Notes

### Competing Interest Statement

The authors have declared no competing interest.

