## Supplementary information for "Diversity and abundance of archaeal amoA genes in the permanent and temporary oxygen minimum zones of Indian Ocean"

Table S1: Physio-chemical parameters and geographical coordinates of sampled stations

| Stations | Depth (m) | DO (µmol. O_2_ L^-1^) | NO_2_+NO_3_ (µmol. N L^-1^) | NH_4_ (µmol. N L^-1^) | Temperature (°C) | Latitude | Longitude |
| --- | --- | --- | --- | --- | --- | --- | --- |
| 1.a | 20 | 167 | 4.8 | 0.12 | 25 | 20° 30' N | 63° 20' E |
| 1.b | 150 | 51.2 | 23.7 | 0.15 | 21 | 20° 30' N | 63° 20' E |
| 1.c | 600 | 3.2 | 3.8 | 0.13 | 11.8 | 20° 30' N | 63° 20' E |
| 1.d | 1500 | 25.9 | 40.1 | 0.02 | 5.2 | 20° 30' N | 63° 20' E |
| 2.a | 20 | 123 | 8.9 | 0.19 | 25 | 15° 10' N | 64° 40' E |
| 2.b | 180 | 19 | 40.2 | 0.15 | 19 | 15° 10' N | 64° 40' E |
| 2.c | 650 | 2.1 | 3.7 | 0.12 | 11.8 | 15° 10' N | 64° 40' E |
| 2.d | 1350 | 27 | 42 | 0.03 | 6.7 | 15° 10' N | 64° 40' E |
| 3.a | 20 | 143 | 14.3 | 0.18 | 25.5 | 11° 25' N | 68° 20' E |
| 3.b | 180 | 7.2 | 7.2 | 0.17 | 19.5 | 11° 25' N | 68° 20' E |
| 3.c | 710 | 2.3 | 2.3 | 0.12 | 11.5 | 11° 25' N | 68° 20' E |
| 3.d | 1620 | 28 | 28 | 0.07 | 4.8 | 11° 25' N | 68° 20' E |
| 4.a | 20 | 213.5 | 9.7 | 0.19 | 25 | 11° 10' N | 86° 15' E |
| 4.b | 70 | 49.2 | 32 | 0.12 | 25.5 | 11° 10' N | 86° 15' E |
| 4.c | 390 | 9.8 | 9.8 | 0.17 | 13.8 | 11° 10' N | 86° 15' E |
| 4.d | 900 | 93.3 | 36.3 | 0.02 | 10.2 | 11° 10' N | 86° 15' E |
| 5.a | 20 | 40 | 1.92 | 0.13 | 25 | 15° 15' N | 87° 25' E |
| 5.b | 65 | 35 | 20.08 | 0.18 | 26 | 15° 15' N | 87° 25' E |
| 5.c | 380 | 8.7 | 35.3 | 0.15 | 13.5 | 15° 15' N | 87° 25' E |
| 5.d | 920 | 81 | 38.2 | 0.07 | 10.7 | 15° 15' N | 87° 25' E |
| 6.a | 20 | 40 | 0.21 | 0.14 | 25 | 18° 55' N | 89° 10' E |
| 6.b | 60 | 37 | 1.2 | 0.11 | 25 | 18° 55' N | 89° 10' E |
| 6.c | 350 | 9.2 | 35 | 0.16 | 13.2 | 18° 55' N | 89° 10' E |
| 6.d | 980 | 61.7 | 38.5 | 0.01 | 10.8 | 18° 55' N | 89° 10' E |

*Sampled stations (1, 2, 3, 4, 5, 6), depth profile (a =surface waters; b = oxycline waters; c = core oxygen minimum waters and d = meso-pelagic waters)

Table S2: amoA gene copy numbers of total archaeal ammonium oxidizers, WCA and WCB ecotypes sampled in both Seas were given.

| **Marginal Seas** | **Ecotypes** | **Station/ depths** | **Subsurface**(gene copies L-1) | **Oxycline**(gene copies L-1) | **Core-OMZ** (gene copies L-1) | **Meso-pelagic** (gene copies L-1) |
| --- | --- | --- | --- | --- | --- | --- |
| **AS** | **amoAt** | station 1 | 4.3x10^6^ | 5.3x10^8^ | 9.2x10^7^ | 1.7x10^7^ |
|  |  | station 2 | 3.8x10^5^ | 4.7x10^7^ | 8.7x10^6^ | 1.4x10^6^ |
|  |  | station 3 | 4.3x10^6^ | 5.3x10^7^ | 3.8x10^6^ | 3.2x10^5^ |
|  |  | **average** | 3.0x10^6^ | 2.1x10^8^ | 3.5x10^7^ | 6.4x 10^6^ |
|  | **WCA** | station 1 | 2.6x10^5^ | 1.3x10^5^ | 1.9x10^7^ | 2.2x10^1^ |
|  |  | station 2 | 4.1x10^4^ | 2.8x10^4^ | 7.2x10^5^ | 3.3x10^2^ |
|  |  | station 3 | 3.8x10^5^ | 2.3X10^5^ | 1.8x10^6^ | 9.4x10^2^ |
|  |  | **average** | 2.7x10^5^ | 1.3x10^5^ | 7.1x10^6^ | 4.3x10^2^ |
|  | **WCB** | station 1 | 0.2x10^2^ | 3.2x10^7^ | 9.9x10^7^ | 6.1x10^6^ |
|  |  | station 2 | 1.2x10^5^ | 1.7x10^5^ | 2.7x10^6^ | 7.6x10^5^ |
|  |  | station 3 | 3.1x10^2^ | 9.1 x10^6^ | 9.3x10^6^ | 3.9x10^4^ |
|  |  | **average** | 4.0x10^4^ | 1.4x10^7^ | 3.7x10^7^ | 2.2x10^6^ |
| **BoB** | **amoAt** | station 4 | 5.0x10^6^ | 3.9x10^7^ | 4.9x10^6^ | 3.8x10^4^ |
|  |  | station 5 | 2.4x10^6^ | 5.5x10^7^ | 3.4x10^5^ | 7.7x10^4^ |
|  |  | station 6 | 7.7x10^6^ | 7.1x10^8^ | 8.7x10^6^ | 9.2x10^4^ |
|  |  | **average** | 4.9x10^6^ | 2.6x10^8^ | 4.6x10^6^ | 6.9x10^4^ |
|  | **WCA** | station 4 | 1.3x10^5^ | 0.2x10^4^ | 1.9x10^5^ | 2.2x10^2^ |
|  |  | station 5 | 2.8x10^3^ | 1.4x10^4^ | 1.8x10^3^ | 0.7X10^1^ |
|  |  | station 6 | 2.7x 10^5^ | 3.1x10^4^ | 4.9x10^4^ | 8.1x10^1^ |
|  |  | **average** | 1.3x10^5^ | 1.5x10^4^ | 8.0x10^4^ | 1.0x10^2^ |
|  | **WCB** | station 4 | 7.1x 10^2^ | 1.3x10^6^ | 4.6x10^6^ | 6.1x10^3^ |
|  |  | station 5 | 2.3x10^1^ | 2.2x10^6^ | 7.2x10^4^ | 7.6x10^4^ |
|  |  | station 6 | 3.3x10^1^ | 1.8x10^7^ | 1.6x10^5^ | 8.8x10^4^ |
|  |  | **average** | 2.5x10^2^ | 7.1x10^6^ | 1.6x10^6^ | 5.6x10^4^ |

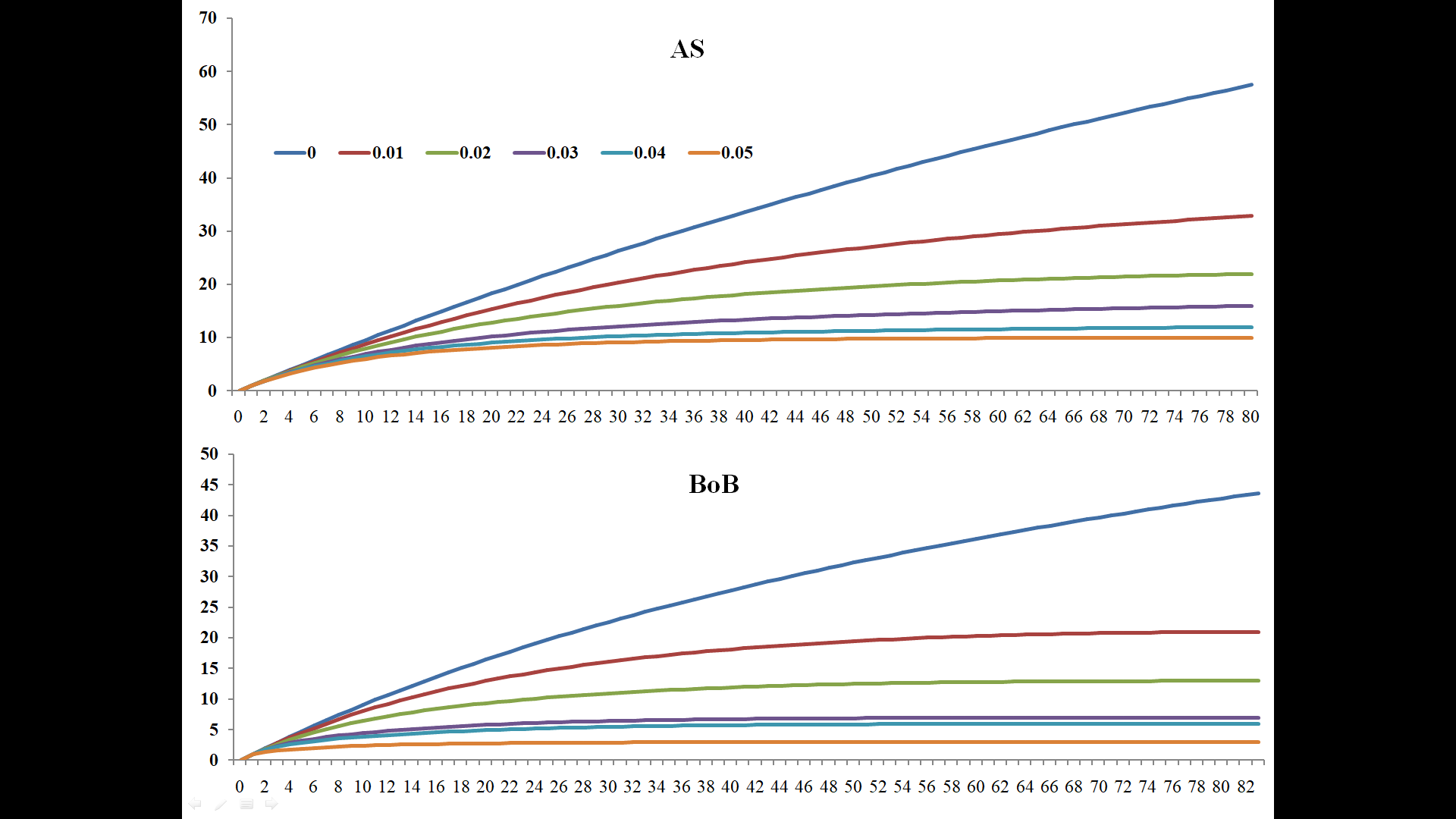

Fig. S1: Rarefaction curve for amoA sequences retrieved from AS and BoB. The X-axis indicates the number of sequences inventoried and Y-axis represents species diversity.

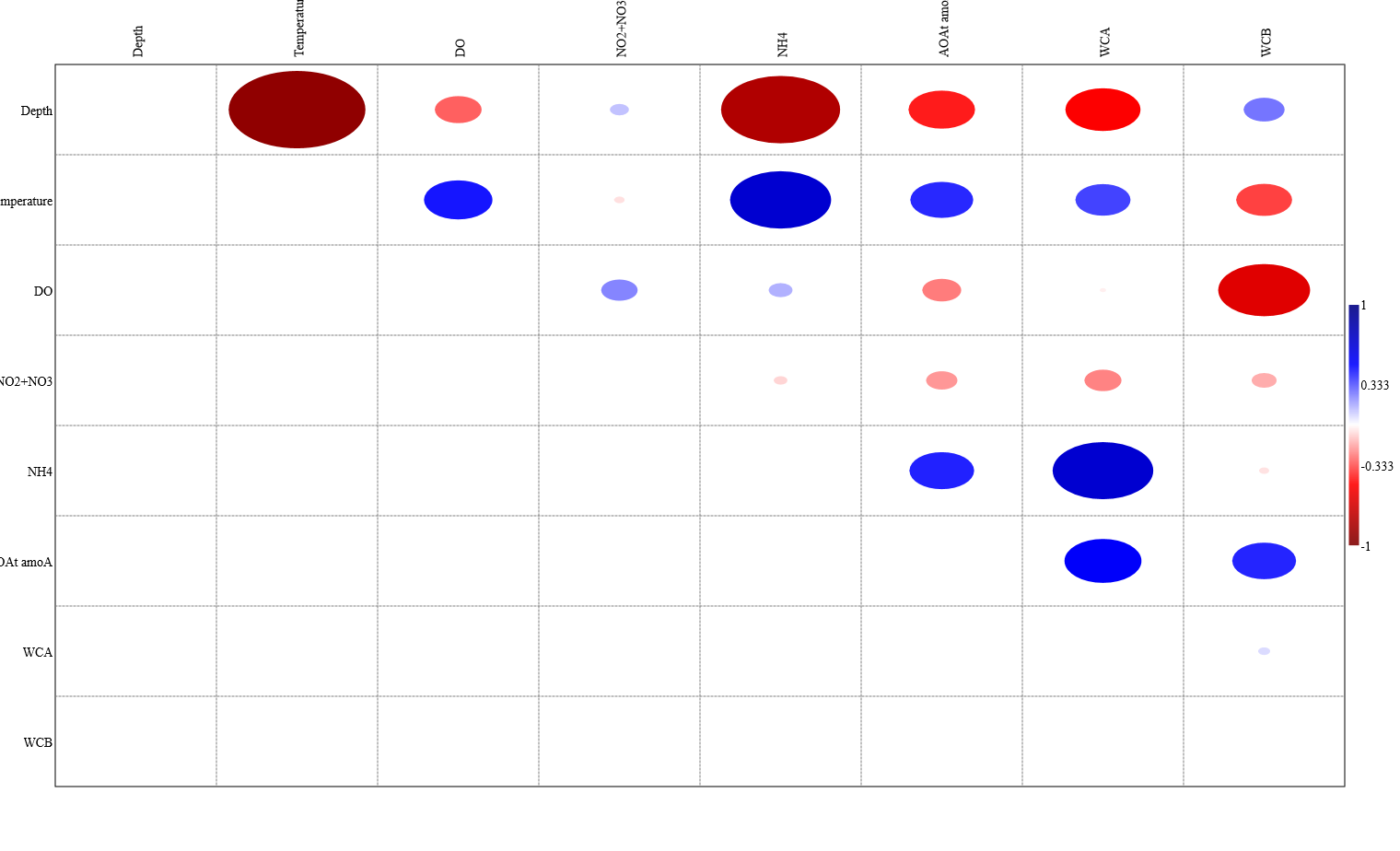

Fig. S2: Correlation oval plot matrix revealing relationships between environmental parameters and gene abundance in AS and BoB

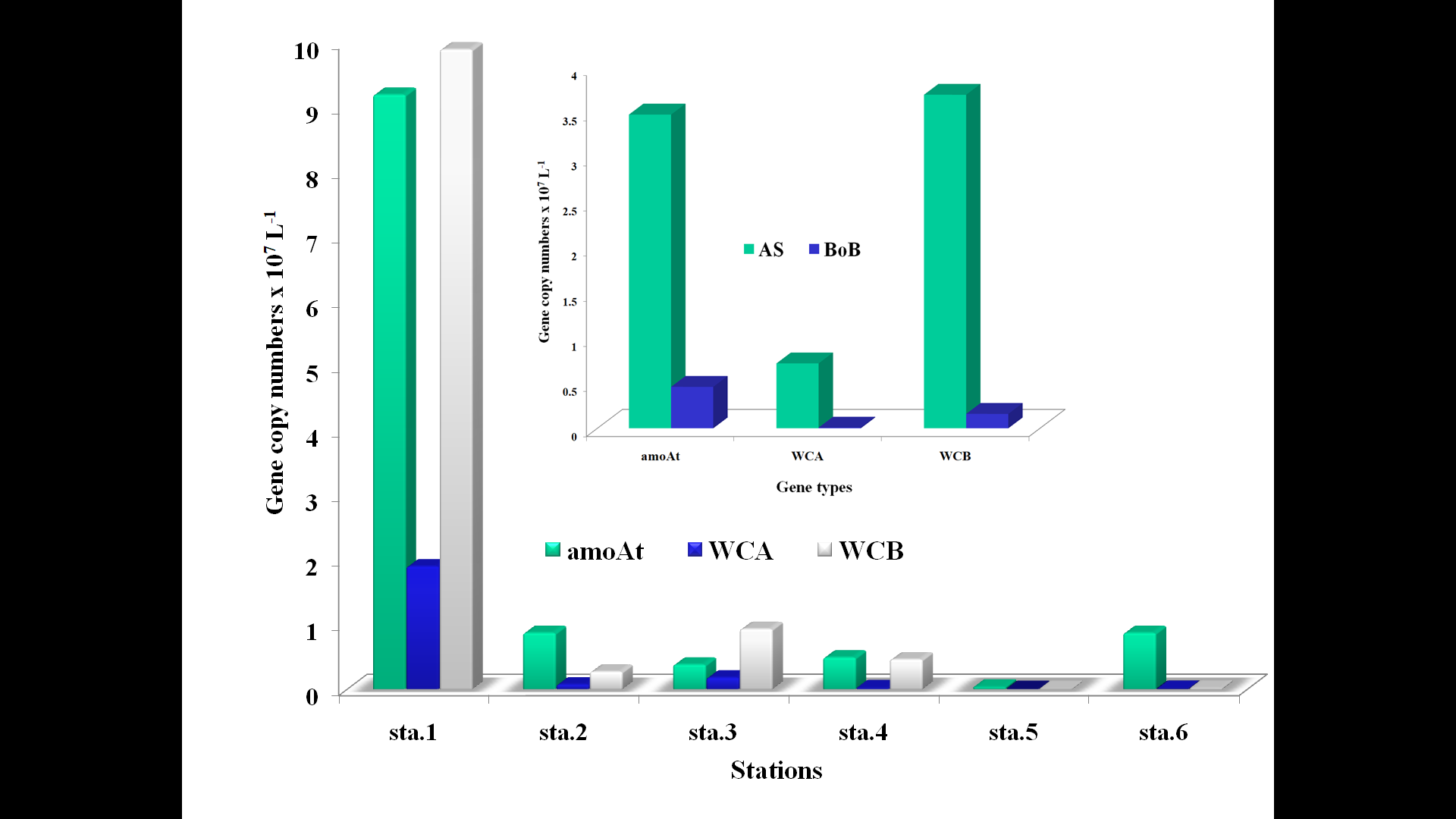

Fig. S3: Comparing abundances of WCA, WCB and amoAt genes in core-OMZs of all sampled stations in AS and BoB. The top picture represents the gene-wise abundance data.

Table S4: BLAST analysis of represented sequences from Alves et al. (2018). Sequences marked red was from the present study used by Alves et al., (2018) for grouping

| GenBank Acc. No | Clade | Iso_Source | Citation |
| --- | --- | --- | --- |
| **JN190773** | **NP-γ-2.2_Incertae** | BoB | Present study |
| **JN190774** | **NP-γ-2.2_Incertae** | BoB | Present study |
| JX524536 | **NP-γ-2.2.4.1_OTU9** | shrimp pond sediment (India) | unpublished |
| GQ863134 | **NP-γ-2.2.4.1_OTU3** | estuarine sediment (Australia) | Abell et al., (2009) |
| FJ227155 | **NP-γ-2.2.4.1_OTU5** | coastal marine sediment (China) | unpublished |
| FJ601588 | **NP-γ-2.2.4.1_OTU8** | tropical marine estuary sediment (India) | Singh et al., (2010) |
| FJ601551 | **NP-γ-2.2.4.1_OTU6** | tropical marine estuary sediment (India) | Singh et al., (2010) |
| **JN190815** | **NP-γ-2.2.4.1_OTU1** | BoB | Present study |
| EU197137 | **NP-γ-2.2.4.1_OTU2** | sediments of Qinghai Lake (China) | unpublished |
| EU025180 | NP-δ-1_OTU2 | Changjiang Estuary sediment (China) | Dang et al., 2008 |
| JF924029 | NP-δ-1_OTU3 | marine surface sediments (China) | Cao et al., (2011) |
| JF924131 | NP-δ-1_OTU10 | marine surface sediments (China) | Cao et al., (2011) |
| **JN190788** | **NP-δ-1_OTU11** | BoB | present study |
| AB373341 | NP-δ-1.Incertae | aquarium biofilter (Japan) | Urakawa et al., (2007) |
| JX283754 | NP-δ-1.Incertae | salt marsh sediment (USA) | Peng et al., (2013) |
| FJ227153 | NP-δ-1_OTU4 | coastal marine sediment (China) | unpublished |
| GQ250745 | NP-δ-1.2_OTU2 | seawater column in the Gulf of Mexico (Mexico) | unpublished |
| **JN190644** | **NP-δ-1.2_OTU1** | AS | Present study |
| AB828780 | NP-ε-2.2_OTU4 | Seawater (USA) | unpublished |
| FJ799203 | NP-ε-2.2_OTU6 | Peruvian oxygen minimum zone (Germany) | Lam et al., (2009) |
| AB625946 | NS-α-3.2.1.1.2 | paddy soil (Japan) | unpublished |
| FR773159 | NS-α-3.2.1.1.1 | The upper layer of garden soil (Austria) | Tourna et al., (2011) |
| **JN190828** | **NS-β-2_OTU3** | BoB | present study |
| AB645413 | NS-β-2_OTU1 | subseafloor sediments off Shimokita Peninsula (Japan) | unpublished |
| FJ601605 | NS-β-2.1_OTU1 | tropical marine estuary sediment (India) | Singh et al., (2010) |
| GQ481090 | NS-β-2.1_OTU3 | Western Amazon Agri soil (Brazil) | Navarrete et al., (2011) |
| **JN190852** | **NS-γ-2.2.3_OTU2** | BoB | present study |
| DQ534701 | NS-γ-2.2.3_OTU3 | Soil (UK) | unpublished |
| **JN190707** | **NS-ε-1_OTU1** | BoB | present study |
| AB828929 | NS-ε-1_OTU2 | Seawater (USA) | unpublished |
| FJ601584 | NS-ε-1_OTU3 | tropical marine estuary sediment (India) | Singh et al., (2010) |
| JN183774 | NS-ε-1_OTU5 | pooled salt-water aquaria (Canada) | Sauder et al., (2011) |
